# Quantifying the Provenance-to-Function Gap in Antidiabetic Peptide Prediction: Homology-Aware Evaluation and the ADP-Hybrid Baseline

**DOI:** 10.64898/2026.09.25.754557

**Authors:** Ariful Islam, Md. Faruk Hosen, Md. Abul Basar, Mohammad Sarwar Hossain Mollah, Muhammad Shahin Uddin

**Author notes:** Corresponding author (Md. Faruk Hosen), (Md. Abul Basar).

## Abstract

Antidiabetic peptides (ADPs) are short bioactive sequences of therapeutic interest, and sequence-based classifiers prioritise experimental candidates. A classifier is useful only if its accuracy transfers to unseen sequences, which depends on how the benchmark was assembled and partitioned. We term the distance between what such a classifier is scored on and the function it is meant to predict the *provenance-to-function gap*, and we quantify it. We re-evaluate the two-layer ADP benchmark of Basith et al. under a protocol that groups homologous peptides rather than splitting them at random. Of the 877 ADPs, 218 attribute to a precursor protein by exact substring containment; within that subset the second-layer label coincides exactly with precursor identity, all 140 human-insulin fragments carrying the type-1 label and all 76 bovine milk-protein fragments the type-2 label, without exception. Accuracy tracks identity to the training set, rising from a Matthews correlation coefficient (MCC) of 0.39–0.43 below 50% identity to 0.92–0.96 between 70% and 90%, and peptide length alone reaches MCC 0.619 on held-out data. Auditing sixteen further peptide benchmarks shows the coupling is not confined to this resource: fragment families share a label more often than chance in eleven of fourteen testable datasets, and not in three, so the property is common rather than universal. Rebuilding the published architecture on identical folds shows its advantage over a single classifier is a function of the split: present under random partitioning, absent once homologues are separated. ADP-Hybrid, one tree-ensemble classifier per layer, reaches MCC 0.843 on layer 1 against a published 0.841 at 21–38 times the inference throughput, and 0.801 against 0.858 on layer 2. We release the protocol and the controls that expose these properties.

## 1. Introduction

Diabetes mellitus is among the most prevalent chronic metabolic disorders worldwide. The International Diabetes Federation estimated 425 million affected adults in 2017 and projected 629 million by 2045, a trajectory driven by population ageing, urbanisation and changes in diet and physical activity [1]. The two principal forms differ in mechanism rather than degree: type-1 diabetes arises from autoimmune destruction of pancreatic *β*-cells and requires insulin replacement, whereas type-2 diabetes is characterised by insulin resistance with progressive secretory failure and is managed through a wider pharmacological repertoire. Because the two conditions present different therapeutic targets, a candidate molecule is of interest not only for whether it is antidiabetic but for which form of the disease it addresses.

Peptides occupy a distinctive position in this therapeutic landscape [2]. They combine the target specificity of biologics with a smaller molecular footprint, and several classes of antidiabetic peptide act through mechanisms that include incretin-receptor agonism and inhibition of enzymes involved in glucose regulation. Candidates arise both from endogenous hormones and their processed fragments and from dietary proteins, and the peptides catalogued in resources such as BioDADPep [3] reflect both origins. This breadth is what makes the space worth searching computationally: the number of sequences that could in principle be assayed vastly exceeds the number that has been.

That gap is the practical motivation for sequence-based prediction. Synthesising a peptide and running an activity assay is slow and expensive enough that exhaustive experimental screening is not a realistic strategy, so a computational filter that ranks candidates before synthesis has become a standard first step, and several predictors are now available for this task [4, 5]. The value of such a filter is entirely a function of how it behaves on sequences that have not yet been characterised — which is to say, on exactly the sequences for which no label exists to check it against.

The usefulness of such a filter depends on a question that reported accuracy alone cannot answer: does performance transfer to peptides unlike those in the training set? This distinction — between *recognising* sequences related to those already seen and *generalising* to sequences that are not — is the central question of a predictor’s utility, because the peptides a screening tool is deployed on are by construction the ones not yet characterised. A model that recognises but does not generalise will report high accuracy on its own benchmark and add nothing to a prospective screen. This is not a given. Peptide-function datasets are assembled from curated activity databases, and those databases are not samples from sequence space. An entry is added because a peptide was reported active, and reports cluster: a single study characterising one precursor protein contributes many overlapping fragments at once. The resulting collection is dense in some regions of sequence space and empty elsewhere, and that density is organised by provenance. Where a class label is defined by therapeutic indication, and different indications were historically investigated in different source proteins, label and provenance become correlated by construction. Under a random train/test split a model can then achieve high accuracy by recognising which protein a fragment came from rather than by learning a determinant of activity.

The general problem is documented across machine-learning-based science [6] and specifically within genomics [7], and community reporting standards now treat the partitioning strategy as a first-class methodological detail [8].

Basith et al. [5] introduced ADP-Fuse, which serves as the reference point throughout this work. Their contribution was substantial: they curated the BioDADPep resource [3] into a two-layer prediction task — separating ADPs from non-ADPs in the first layer, then assigning an ADP to type-1 or type-2 diabetes in the second — conducted a broad comparison of 18 descriptor families against eight classifier families, and released a public web server. The datasets and the two-layer formulation used here are theirs, employed without modification.

Figure 1 summarises the structure of the study. Three questions motivate it. First, what is the structure of this benchmark, and what can a classifier exploit in it? Second, how does accuracy distribute across sequences of differing similarity to the training data, which requires a homology-aware partition rather than a random one? Third, how much model complexity does the achievable accuracy actually require? The principal contributions are as follows.

**Figure 1:**
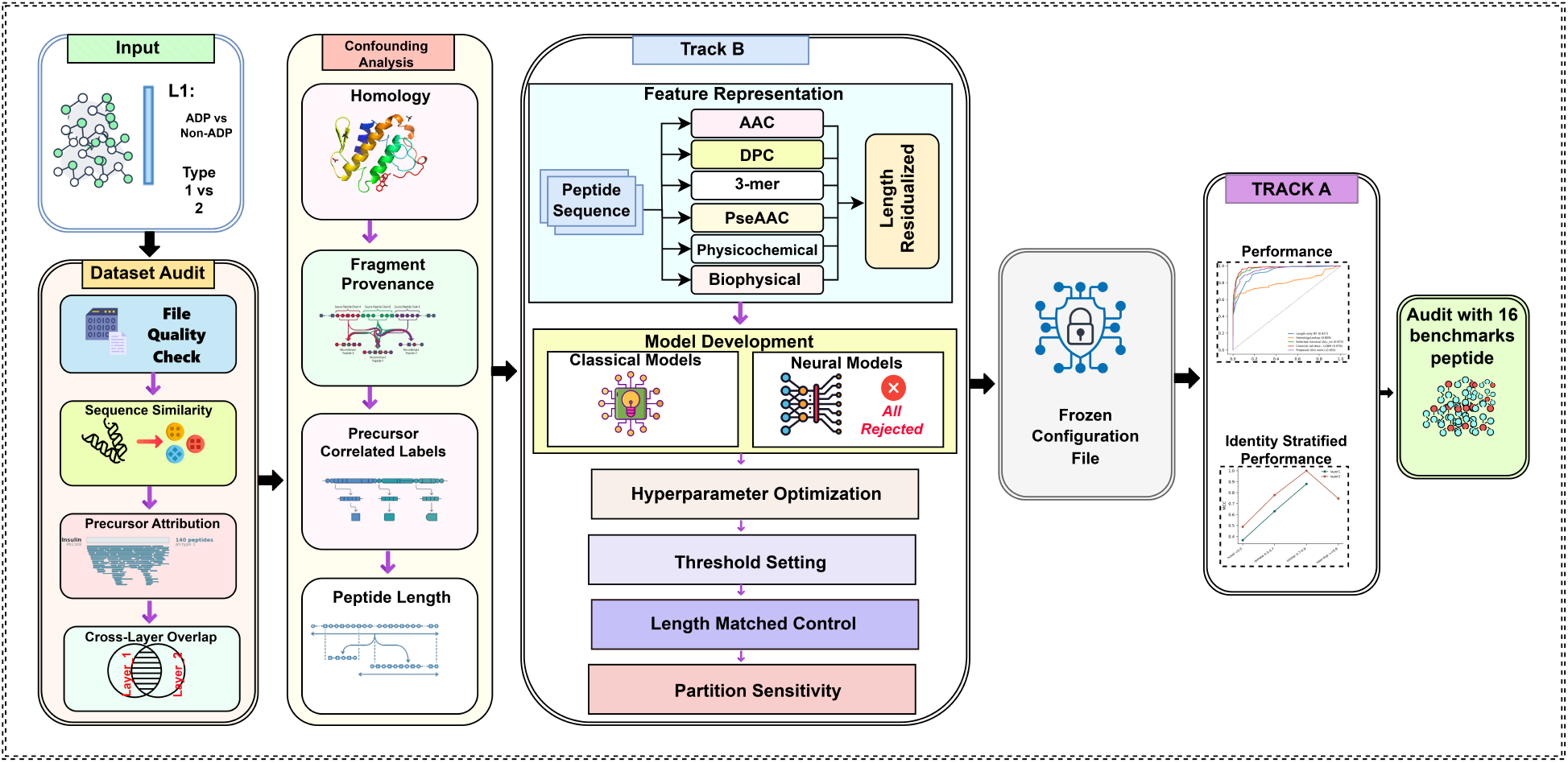
Structure of the study. Four audits are conducted on the training partition alone; each yields a property of the benchmark that bears on how a reported accuracy should be read. Those properties motivate a corrected evaluation protocol, under which a single locked model is scored. Every quantity shown is reported with its provenance in Section 4.

- The second-layer class label coincides with precursor-protein identity for every peptide that could be attributed — 218 peptides across five precursors, with no exceptions (Section 4.2).
- Predictive accuracy tracks sequence identity to the training set monotonically, and 13.4% of the first-layer independent test set forms a single-class stratum that cannot be scored (Section 4.8).
- A homology- and length-aware evaluation protocol is provided, including a length-matched control whose own sanity check is reported (Section 3.4).
- ADP-Hybrid, a single tree-ensemble classifier per layer, reaches first-layer accuracy that the available evidence does not separate from the published 43-model ensemble, at 21–38*×* the inference throughput (Sections 4.8 and 4.9).
- A bounded negative result on architecture: six neural architectures, including frozen protein language model representations, did not exceed tuned descriptors under matched conditions (Section 4.5).

## 2. Related work

AntiDMPpred [4] applied random forests to composition-derived features, reporting an accuracy of 0.771 and an MCC of 0.544 on its own data. ADP-Fuse [5] extended the task to two layers and introduced a multiview stacking framework in which single-feature models over 18 descriptor sets supply predicted probabilities and class labels to a meta-classifier, reporting cross-validated MCC values of 0.886 and 0.917 and independent-test values of 0.841 and 0.858 for the two layers respectively.

The multiview stacking design of ADP-Fuse follows a lineage of peptide-function metapredictors in which many single-descriptor models supply probabilities to a second-stage learner. mAHTPred [9] established the pattern for antihypertensive peptides, combining several composition and physicochemical encodings under a meta-classifier, and the design has since been carried across a range of peptide activities. These frameworks consistently report accuracy gains over their single-model components, and those gains are real under the protocol used to measure them. What is less often reported alongside them is the partitioning under which they were obtained, or the accuracy of a trivial covariate baseline on the same split — the two quantities that determine whether a gain reflects a better representation or a more redundant benchmark. The present work supplies both for one such framework.

That homology between partitions inflates reported accuracy in peptide bioactivity prediction is already established, and the present study should be read as building on that work rather than introducing the concern. Fernández-Díaz et al. [10] surveyed 18 peptide bioactivity datasets and found that 13 contained at least 10% of training peptides similar to test peptides; only five applied any homology-based correction, and those used 80–90% identity thresholds permissive enough to leave substantial leakage. Re-partitioning by homology reduced reported performance by up to 50% on some tasks. Sidorczuk et al. [11] showed independently that antimicrobial peptide benchmarks are biased by how the negative class is sampled, so that reported accuracy partly reflects the sampling procedure rather than the biology. Both studies establish the general phenomenon across families of peptide predictors. What neither addresses, and what this study contributes, is the curation mechanism that generates the redundancy in a specific benchmark: not that homologues straddle the split, but why they do, and what the class label therefore encodes.

The concern is not confined to peptide work: Luo et al. [12] found that 85 of 93 test proteins in a widely used DNA-binding benchmark shared more than 40% identity with its training set, and that eleven predictors fell from roughly 0.85 to roughly 0.1 MCC once that redundancy was removed.

Grouping homologous sequences before partitioning is routine practice in structure prediction and remote-homology benchmarks, and efficient tools exist for it at scale [13, 14], but it is applied inconsistently in peptide-function work [10], where redundancy reduction is sometimes applied to a single class in order to preserve sample size. Against that background the contribution of the present work is narrower than a general demonstration and correspondingly more specific: to quantify, for one widely used two-layer peptide benchmark, how much of the reported accuracy is attributable to sequence relatedness between partitions, and to identify the mechanism — precursor-correlated labelling — by which that relatedness arises in this resource.

## 3. Materials and Methods

### 3.1. Dataset description

The two-layer dataset distributed with ADP-Fuse [5], derived from BioDADPep [3], is used without modification. The first layer discriminates ADPs from non-ADPs; the second assigns an ADP to type-1 or type-2 diabetes. Composition is given in Table 1. All four distributed files parse cleanly, contain no non-standard residues, no duplicate identifiers and no within-file duplicate sequences, and their counts otherwise reconcile with the published description; one discrepancy in the training fraction is reported in Section 4.1.

**Table 1:** Composition of the two-layer benchmark as distributed. For layer 1, class 1 is ADP and class 2 non-ADP; for layer 2, class 1 is type-1 and class 2 type-2, the two diabetes types rather than a presence/absence contrast. Class-mean peptide lengths are reported because their separation motivates the length-matched control described in Section 3.4.

| Layer | Split | $n$ | Class 1 | Class 2 | Mean len. (1) | Mean len. (2) |
| --- | --- | --- | --- | --- | --- | --- |
| 1 | Training | 1,316 | 658 | 658 | 13.98 | 25.73 |
| 1 | Independent | 960 | 219 | 741 | 13.73 | 26.55 |
| 2 | Training | 700 | 500 | 200 | 15.42 | 9.50 |
| 2 | Independent | 177 | 155 | 22 | 15.55 | 8.41 |

### 3.2. Sequence identity, clustering and precursor attribution

Pairwise sequence identity is computed by semi-global alignment [15] with free end gaps (match +1, mismatch 0, gap opening − 1, gap extension − 0.5) using Biopython [16], taking the shorter of the two sequences as the denominator, which is the convention adopted by CD-HIT [13, 17]. For two sequences *a* and *b* with an optimal alignment containing *n*_id_ identical aligned residues, identity is defined as

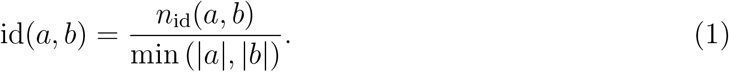

Two clustering algorithms are computed. The primary is a greedy incremental procedure equivalent to CD-HIT: sequences are processed longest-first, and a sequence joins an existing cluster only if its identity to that cluster’s representative meets the threshold. Single-linkage clustering is computed as a conservative bound; because it chains, it collapses the data considerably more aggressively at low thresholds, and greedy figures are therefore reported throughout unless stated otherwise.

Precursor attribution tests each peptide for exact substring containment within a curated panel of candidate precursor proteins retrieved from UniProt [18], with nine additional proteins included as negative controls. No feature derived from precursor identity enters any predictive model at any stage.

### 3.3. Feature representation

Candidate descriptor blocks comprise amino-acid composition (AAC, 20D), dipeptide composition (DPC, 400D), 3-mer composition restricted to the 400 most frequent 3-mers observed in the training split (400D, rather than the full 20^3^ = 8000 tripeptides, most of which never occur in peptides of this length), composition–transition–distribution descriptors (CTD, 273D), pseudo amino-acid composition [19] (25D), a physicochemical block (18D) constructed from Kyte–Doolittle hydropathy [20] and the Eisenberg hydrophobic moment [21], and a biophysical block (13D) comprising Chou–Fasman conformational propensities [22], net charge at pH 7.4, isoelectric point, and the aliphatic [23] and Boman [24] indices.

For a peptide *s* of length *L*, amino-acid composition is

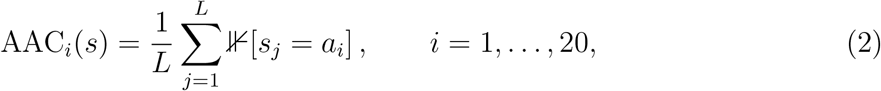

and dipeptide composition is

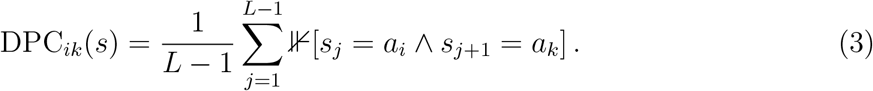

Net charge at a given pH is obtained from the Henderson–Hasselbalch relation over ionisable groups,

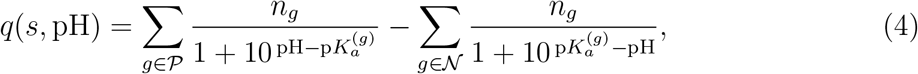

Where *P* and *N* denote the sets of positively and negatively ionisable groups and *n*_*g*_ their counts; the isoelectric point is the pH at which *q*(*s*, pH) = 0, obtained by bisection. The Eisenberg hydrophobic moment at rotation angle *δ* is

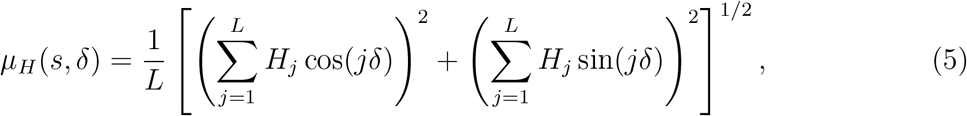

with *H*_*j*_ the hydropathy of residue *j* and *δ* = 100^*°*^ for an *α*-helix.

Candidate classifiers are LightGBM [25], XGBoost [26], histogram gradient boosting [27] and CatBoost [28]. For the architecture comparison we additionally evaluated a Transformer [29] trained from scratch, a convolutional–Transformer hybrid, three variants built on frozen ESM-2 650M representations [30], and convolutional, bidirectional LSTM [31], hybrid convolutional–recurrent and attention-augmented recurrent baselines.

### 3.4. Validation protocol

Two evaluation tracks are reported throughout. *Track A* reproduces the original protocol: a model is trained on the training file and evaluated once on the independent file. *Track B* is cluster-grouped cross-validation conducted entirely within the training file at 50%, 70% and 90% identity, with 70% treated as primary. All cross-validation uses ten cluster-disjoint folds; paired tests over folds therefore have *n* = 10, and the smallest attainable two-sided Wilcoxon signed-rank [32] *p*-value is 2*/*2^10^ = 0.00195. All model development, descriptor-block selection, hyperparameter search and threshold setting occur within Track B. Inner validation folds used for early stopping are also split by cluster, so that early stopping never observes a homologue of the data against which it is scored.

A *length-matched control* pairs each positive peptide with an unused negative of exactly the same length, retaining matched pairs within the same fold by merging pair links and cluster links under a union–find structure [33] before the folds are cut. The control is validated by training the same random forest on peptide length alone over the matched subset: if the matching is correct that model has no information and must score MCC 0.000, which is the check reported below. Two design details proved necessary, and each was identified by the control failing its own sanity check: a tolerance of *±*1 residue was insufficient, as gradient boosting on length alone still attained MCC 0.43, and splitting matched pairs across folds drove the length-only baseline to MCC − 0.45 rather than zero. With both corrected, length scores exactly MCC 0.000, which constitutes the evidence that the control functions as intended.

Model comparison uses the Matthews correlation coefficient [34] as the primary metric, this being appropriate for imbalanced binary problems [35]. Independent-test metrics carry stratified percentile bootstrap 95% confidence intervals computed over 2,000 resamples [36]. Within-study model pairs are compared by paired bootstrap on identical resamples and by DeLong’s test for correlated areas under the receiver operating characteristic curve [37], with Holm–Bonferroni correction [38] applied within each comparison family (ablation, head-to-head) rather than across the study as a whole. Two decision thresholds are declared in advance: the default 0.5, and a threshold corrected analytically for the shift in class prior between training and deployment. Writing *π*_tr_ and *π*_te_ for the training and deployment positive-class priors, the corrected threshold *τ* ^*′*^ satisfies

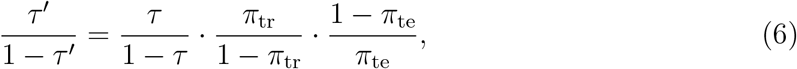

with *π*_te_ taken from the class counts published by Basith et al. [5] rather than measured on the test files.

We refer to the configuration selected by this procedure as ADP-Hybrid. On layer 1 it is a random forest [39] over the length-residualised concatenation of all descriptor blocks (843 features); on layer 2, gradient boosting over the same representation (294 features after the layer-2 blocks are assembled). Neither layer retains any neural component: the search removed them when given the choice. Selected blocks, hyperparameters, decision thresholds and ensemble sizes for both layers are listed in Supplementary Table S6, and the frozen configuration file is released with the code. This choice keeps the independent files sealed during threshold setting, at a cost worth stating plainly: the corrected threshold is optimal for the deployment prior as published, not for the empirical prior of the test set. Where the two differ the threshold is therefore slightly mis-set relative to what a post-hoc fit would give, which is the price of not looking.

### 3.5. Hyperparameter optimisation

Hyperparameters were optimised with Optuna [40] using a tree-structured Parzen estimator sampler, with mean MCC over the 70%-identity cluster-grouped folds as the objective. Two rounds were conducted: an initial search of 60 trials per layer, followed by an expanded search of up to 250 trials restricted to the best-performing descriptor families, adding Cat-Boost as a fourth classifier family, the biophysical block described in Section 3.3, and pairwise product and ratio terms among the ten features ranked highest by mean absolute SHAP value [41]. Both layers ran the full 250 trials. On the first layer the best value was reached at trial 38 and was not improved by any of the 211 trials that followed; an earlier version of this analysis stopped that search at 133 trials on a plateau criterion, and completing it returned an identical configuration and an identical objective value, so nothing downstream of the search changes. The second layer behaved differently: its best value was first reached at trial 245 of 250, so that search had not plateaued when the budget ran out and a longer run might improve the second-layer configuration. We report this rather than extend the budget, because a better configuration would require re-locking and a further evaluation of an independent set that has already been opened twice. Residue-level attribution for the interpretability analysis used integrated gradients [42]. The selected configuration, its decision threshold and its ensemble size were frozen to a configuration file before the independent test files were opened for scoring. The complete search spaces for every descriptor block, classifier family and neural component are given in Supplementary Table S31.

### 3.6. Software environment

All analysis was carried out under Python 3.11.15 on macOS (Apple silicon), with NumPy 1.26.4 [43], pandas 2.2.2 [44], SciPy 1.17.1 [45], scikit-learn 1.8.0 [27], LightGBM 4.7.0, XG-Boost 2.0.3, CatBoost 1.2.10, PyTorch 2.12.0 [46], Biopython 1.86, Optuna 4.8.0, SHAP 0.51.0 and Matplotlib 3.8.4 [47]. Neural models were trained on the Metal Performance Shaders backend. Random seeds are fixed in every script; the seed-stability figures in Section 4 come from five independent end-to-end retrainings under different seeds.

### 3.7. Evaluation metrics

Given the confusion-matrix counts TP, TN, FP and FN, the reported metrics are defined as follows:

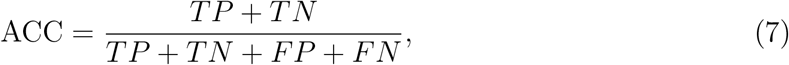

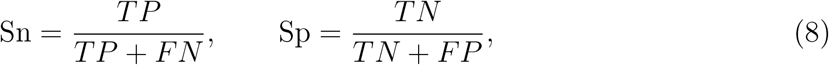

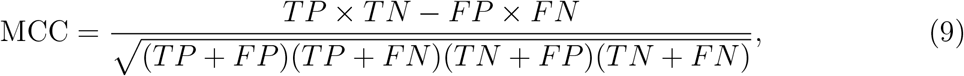

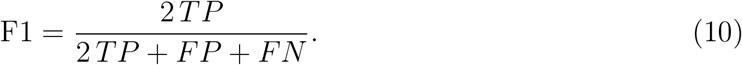

The area under the receiver operating characteristic curve (AUC) and the area under the precision–recall curve are additionally reported as threshold-independent measures.

## 4. Results

### 4.1. Audit of the distributed files

All four files parse without error, contain only the twenty standard residues, and carry no duplicate identifiers or within-file duplicate sequences (Supplementary Figure S1). Two points arise on reconciling them with the published description. The original manuscript states that 80% of the ADPs were used for first-layer training, whereas the files give 658*/*877 = 75.0%; no peptides are missing, so the discrepancy is one of reporting, but a reader reconstructing the splits from the text alone would arrive at different counts. Second, the layer-2 files partition the same 877 peptides differently from layer 1, with consequences taken up below.

### 4.2. The second-layer label coincides with precursor identity

Attributing the 877 ADPs to precursor proteins by exact substring containment yields a complete separation by precursor within the attributable subset (Table 2). All 140 human-insulin fragments carry the type-1 label and none the type-2 label; all 76 bovine milk-protein fragments carry the type-2 label and none the type-1 label. Across the 218 attributable peptides there are no exceptions. That milk proteins are a documented source of DPP-IV-inhibitory antidiabetic peptides [48, 49, 50] makes the coincidence unsurprising as biology, and all the more consequential as a label. Nine control proteins — pro-glucagon, gastric inhibitory polypeptide, insulin-like growth factors 1 and 2, pancreatic polypeptide, vasopressin-neurophysin, albumin, haemoglobin and myoglobin — contributed no fragments, indicating that the attribution is specific rather than an artefact of substring search.

**Table 2:** Attribution of the 877 ADPs to precursor proteins by exact substring containment against sequences retrieved and verified from UniProt [18]. Within the attributable subset the second-layer class label is fully determined by precursor identity.

| Precursor protein (UniProt) | Length | Fragments | % of set | Type-1 | Type-2 |
| --- | --- | --- | --- | --- | --- |
| Human insulin (P01308) | 110 | 140 | 16.0 | 140 | 0 |
| Bovine β-casein (P02666) | 224 | 61 | 7.0 | 0 | 61 |
| Bovine β-lactoglobulin (P02754) | 178 | 12 | 1.4 | 0 | 12 |
| Bovine α-S1-casein (P02662) | 214 | 3 | 0.3 | 0 | 3 |
| Human IAPP / amylin (P10997) | 89 | 2 | 0.2 | 2 | 0 |
| Attributable total |  | 218 | 24.9 | 142 | 76 |

It is important to state precisely what this establishes. The separation is exact within the 24.9% of the set that could be attributed. The remaining 659 peptides are unattributed, and it cannot be verified that they behave in the same way. The attributed subset is moreover not a random sample of the whole: it is enriched for type-2 peptides relative to the full set (65.1% type-1 against 74.7%), as would be expected when one precursor family yields short and compositionally distinctive fragments. Exact substring containment further misses homologues, sequence variants and post-curation edits, so 24.9% is a lower bound on the precursor-derived fraction. The defensible statement is therefore that *for every peptide whose precursor could be identified, the second-layer label is predictable from that precursor alone*; whether the unattributed remainder shares this structure remains untested.

This structure additionally accounts for the compositional difference between the two second-layer classes without recourse to mechanism. Bovine *β*-casein is proline-rich and yields short peptides, and correspondingly, in the second-layer training split, the type-2 ADPs average 9.5 residues at 18.2% proline against 15.4 residues and 4.7% proline for the type-1 peptides. The same contrast holds across both splits combined (9.4 residues at 17.8% proline against 15.5 at 4.7%).

### 4.3. Redundancy structure differs between the classes

The original study applied CD-HIT at a 0.6 threshold to the negative class and, as stated in its methods, did not apply it to the positive class in order to preserve sample size. The resulting asymmetry is directly observable (Supplementary Table S34): 41.4% of ADPs are proper fragments of another ADP, against 0.0% of non-ADPs, and 54.3% of independent-test ADPs stand in a containment relation with a training ADP — one sequence is a subsequence of the other, so their identity is 1.000 under the convention of Equation (1) — against 0.0% of test non-ADPs. Of those 119 peptides, 85 are fragments of a training peptide and 73 contain one, the two sets overlapping. The one-directional figure, test peptides that are proper fragments of a training peptide, is 38.8%. Of the 107 multi-member fragment families identified among the ADPs, 59 have members on both sides of the train/test split; the twelve largest are listed in Supplementary Table S3, and all twelve straddle it. Human preproinsulin is covered across all 110 of its residues by overlapping peptides, to a maximum depth of 29.

The consequences of that asymmetry for evaluation are shown in Figure 3, which places the identity distribution of each class alongside the effective sample size surviving clustering and the best accuracy attainable as the partition is progressively tightened.

The three panels describe one mechanism from three directions. The positive and negative identity distributions in panel (A) barely overlap, so the two classes do not present the same problem to a classifier. Panel (B) shows what that costs statistically: at a 0.5 threshold the second layer retains 31.3% of its nominal size, so the effective sample is roughly a third of the 700 peptides the file appears to contain. Panel (C) is the consequence for reported accuracy. It plots the best MCC over *all* evaluated models at each threshold, which is not the same quantity as any single row of Table 3: the best model at 0.7 identity scores 0.811, whereas the concatenated descriptor block of Table 3 scores 0.804 there. On that best-of-all-models measure accuracy falls as the partition tightens, monotonically on the second layer and with one reversal on the first, where the 0.5-identity partition recovers to 0.817 from 0.811 at 0.7, for the reason given in Section 4.5. Together these indicate that a figure quoted under a random split describes performance on a largely homologous test set rather than on novel sequences.

**Table 3:**
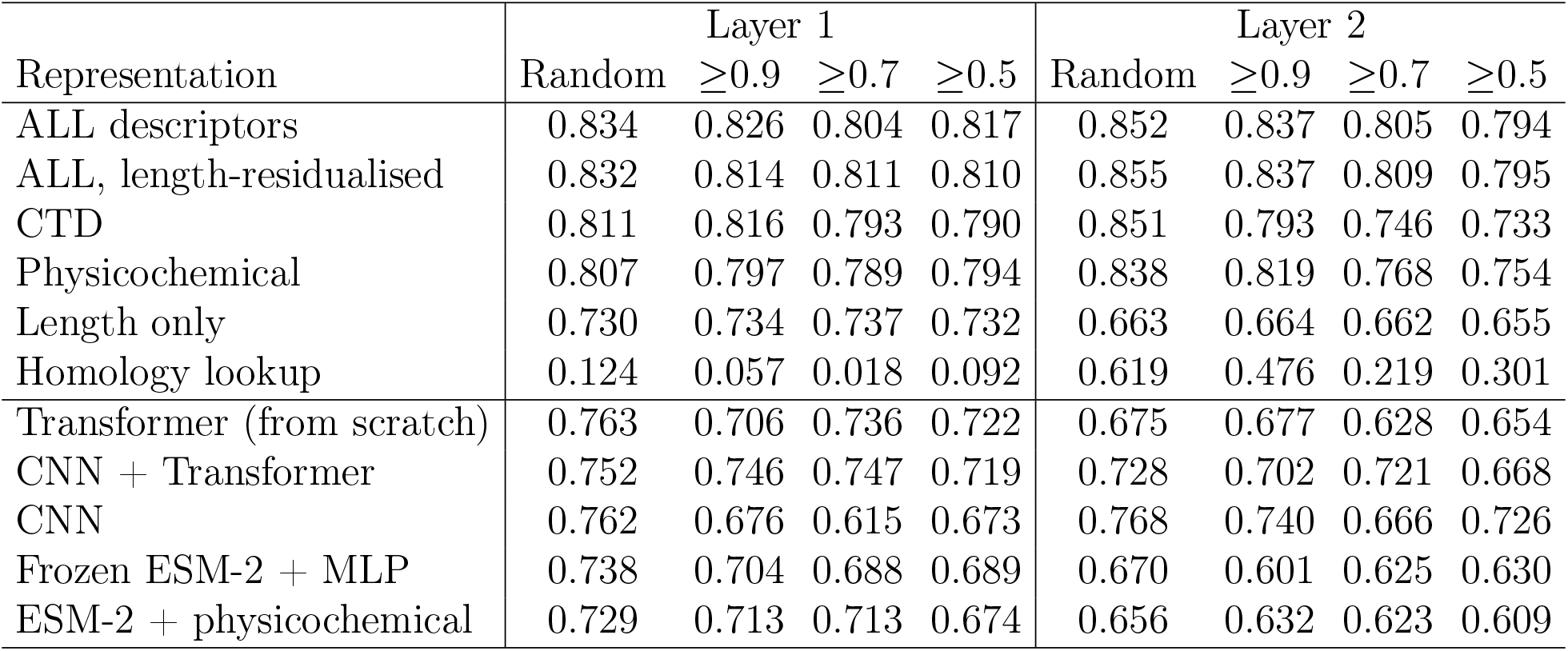
Cross-validated MCC by partition. `ALL` denotes the concatenation of all descriptor blocks and `ALL_noLen` its length-residualised variant. The homology lookup is a feature-free baseline: for each held-out peptide it finds the single most similar training peptide under Equation (1) and returns that peptide’s label, so it uses no descriptor and measures only what nearest-neighbour identity supplies.

A related consequence concerns the two-layer cascade. The 877 ADPs were partitioned independently for each layer, so while each layer’s independent evaluation is internally valid — and the original work reports them in exactly that way — 176 of the 219 first-layer test ADPs (80.4%) also appear in the second layer’s training set. Only 43 peptides are unseen by both layers, which limits what can be established about the cascade evaluated end to end. This is a structural property of independently partitioned multi-stage frameworks in general rather than anything specific to this dataset.

### 4.4. Contribution of peptide length

ADPs average 14.0 residues and non-ADPs 25.7 (Table 1), and the distributions are separated in both the training and the independent split (Figure 2). A random forest supplied with peptide length as its only feature attains MCC 0.729 in cross-validation and MCC 0.619 (AUC 0.927) on the held-out set, which is 74% of the published independent-test MCC obtained from a single integer. Under the length-matched control the length-only baseline correctly collapses to MCC 0.000, while descriptor models retain MCC 0.437 on layer 1 and 0.356 on layer 2 under cluster-grouped validation. Sequence signal beyond length therefore exists and is substantial, at approximately half the magnitude of the unadjusted figures.

**Figure 2:**
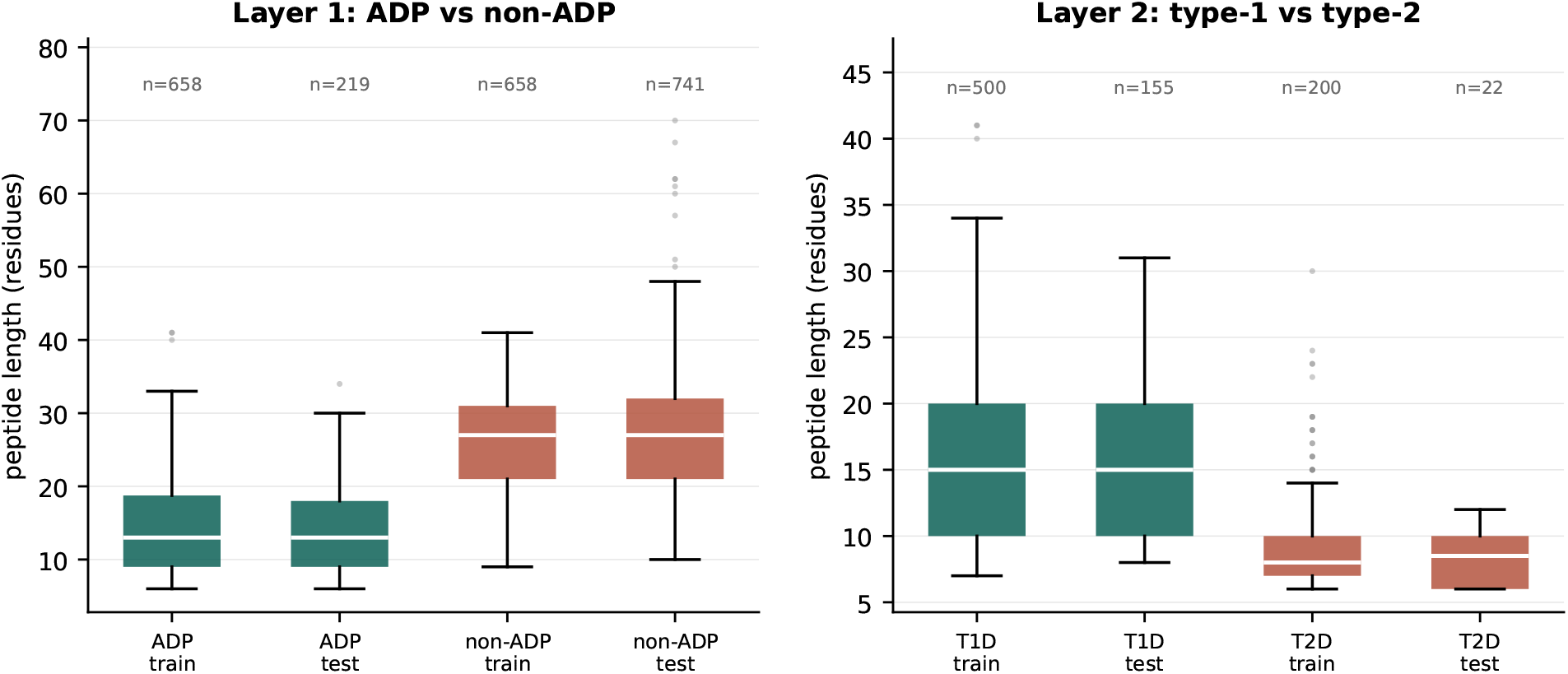
Peptide-length distributions by class and split. The classes are separated by roughly twelve residues in the first layer and six in the second, and the separation is reproduced in the independent split, so a model has access to it at test time. This is what the length-matched control of Section 3.4 removes, and what the length-only baseline of Section 4.4 quantifies. Boxes span the interquartile range, whiskers extend to 1.5*×* IQR, and points beyond are drawn individually.

**Figure 3:**
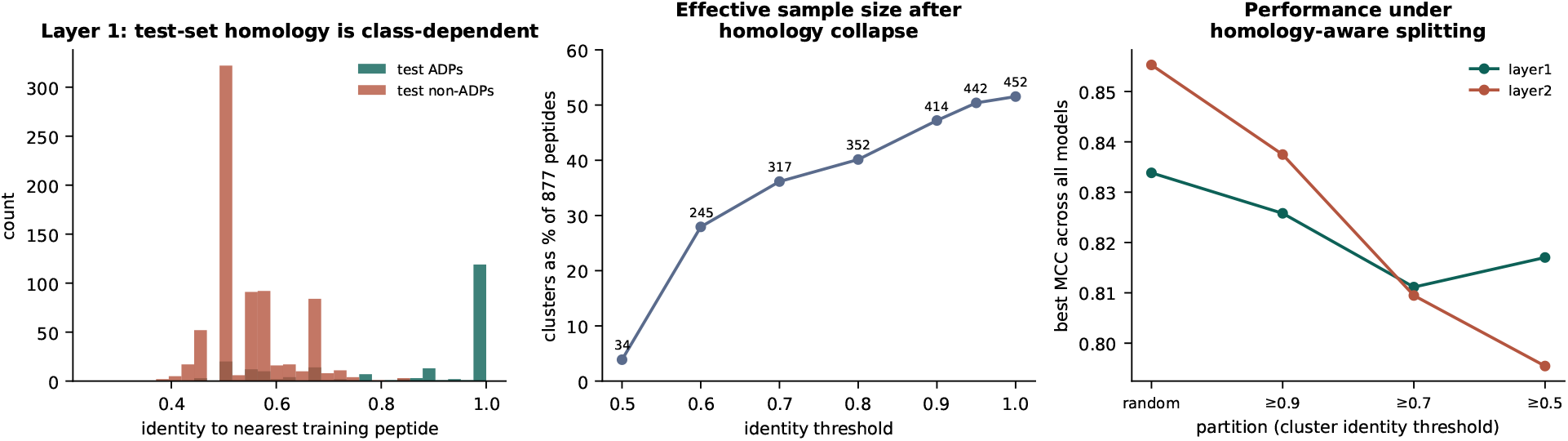
Homology structure of the benchmark. (A) Nearest-neighbour identity from each first-layer independent-test peptide to the training set, separately by class. (B) Clusters obtained from the 877 ADPs as a function of identity threshold, as a percentage of the nominal sample size. (C) Best Matthews correlation coefficient across all evaluated models as the partition becomes progressively homology-aware.

### 4.5. Cross-validated performance

Table 3 reports cross-validated MCC by partition for the strongest descriptor blocks and the strongest neural architectures. Two observations hold across both layers. Performance under the greedy cluster-grouped protocol is below the random-split figure for the descriptor blocks, which is the expected behaviour when related sequences are prevented from straddling a partition. Two qualifications are needed and neither is cosmetic. The decline is not monotonic in the threshold: on layer 1 the concatenated block scores 0.817 at 0.5 identity against 0.804 at 0.7, because greedy clustering at a lower threshold merges more sequences into fewer, larger clusters, so the difficulty of the split and the effective sample size change together and need not move performance in the same direction. Nor is the fall universal: CTD on layer 1 is 0.005 higher at 0.9-identity clustering than under a random split, a difference well inside fold-to-fold variation. The length-only baseline is flat across all four protocols on both layers, which is the result one should expect — peptide length carries no information about homology, so partitioning by homology cannot degrade it. That the descriptor blocks fall while the length baseline does not is itself evidence that what the descriptors lose under cluster-grouping is relatedness rather than signal. Second, no neural architecture exceeded tuned descriptors in any cell.

The ordering in Supplementary Figure S7 is stable across all four partition protocols: descriptor blocks lead, neural architectures follow, and the feature-free homology lookup trails on the first layer while performing unexpectedly well on the second. That last point is diagnostic rather than encouraging — a baseline predicting from training-set identity alone reaches MCC 0.619 on the second layer under a random split, which is a property of the partition and not of the peptides. Complete per-cell values for every descriptor and classifier combination are given in Supplementary Tables S11 and S12, and for every neural architecture in Supplementary Tables S15 and S16.

#### 4.5.1. Architecture search and ablation

The neural hyperparameter search set `use_encoder = False` independently on both layers; given free choice, the optimiser removed the Transformer encoder. Component ablation conducted under the primary protocol agrees. Removing both the pretrained language model and the convolutional motif extractor *improved* first-layer MCC by 0.079 (Wilcoxon signed-rank *p* = 0.0059, Holm-adjusted *q* = 0.041), the only effect in either layer that survives multiplicity correction. Removing the encoder or the convolutional branch alone had no detectable effect, and the one component whose removal cost performance, attention pooling, did not survive correction.

Table 4 reports every comparison; the sign convention there is (full − ablated), so the improvement above appears as Δ = − 0.079.

**Table 4:** Component ablation under the primary 70%-identity cluster-grouped partition. Δ is (full model − ablated model), so a positive value means that removing the component reduced performance. *p* is the Wilcoxon signed-rank test over folds, *q* its Holm-adjusted value within each layer’s family of seven comparisons, and *d*_*z*_ the paired effect size. Only one row in the table is significant after correction, and it avours the reduced model.

| Configuration | Layer 1 |  |  |  |  | Layer 2 |  |  |  |  |
| --- | --- | --- | --- | --- | --- | --- | --- | --- | --- | --- |
| | MCC | SD | $\Delta$ | $q$ | $d_z$ | MCC | SD | $\Delta$ | $q$ | $d_z$ |
| Proposed (full) | 0.660 | 0.086 | — | — | — | 0.651 | 0.133 | — | — | — |
| – motif convolution | 0.656 | 0.094 | +0.004 | 1.000 | +0.05 | 0.602 | 0.166 | +0.050 | 0.654 | +0.46 |
| – Transformer encoder | 0.663 | 0.064 | −0.003 | 1.000 | −0.07 | 0.584 | 0.200 | +0.067 | 0.654 | +0.49 |
| – attention pooling | 0.603 | 0.112 | +0.058 | 0.387 | +0.57 | 0.529 | 0.163 | +0.122 | 0.342 | +0.74 |
| – physicochemical block | 0.652 | 0.090 | +0.008 | 1.000 | +0.12 | 0.553 | 0.198 | +0.098 | 0.342 | +0.59 |
| – pretrained pLM | 0.699 | 0.049 | −0.039 | 1.000 | −0.43 | 0.665 | 0.167 | −0.014 | 1.000 | −0.09 |
| – **pLM – conv** | **0.739** | 0.058 | **−0.079** | **0.041** | −0.96 | 0.681 | 0.105 | −0.030 | 1.000 | −0.27 |
| encoder only (minimal) | 0.652 | 0.097 | +0.008 | 1.000 | +0.11 | 0.614 | 0.154 | +0.037 | 0.654 | +0.35 |

The pattern is consistent across the table: the two components a reader would expect to matter most are the two whose removal helps. On the second layer no effect survives correction at all, and the fold-to-fold standard deviations (0.11–0.20) exceed most of the differences being compared, which is the honest summary of that layer.

The fold-to-fold standard deviations overlap for most pairs in the table, which is why only one comparison survives correction; Supplementary Figure S5 plots the same ablation with those deviations as error bars, and the ablation under all three partition protocols is given in Supplementary Tables S20 and S21.

The resulting claim is stated narrowly: *under matched partitions and search budgets, none of the architectures tested exceeded tuned descriptors on this dataset*. This is not evidence that attention-based models are unsuitable for peptide-function prediction in general. ESM-2 was used in frozen form only; fine-tuning was not attempted, and the neural search budget was 60 trials per layer. What the data regime suggests is nevertheless unsurprising: at 6–41 residues, a few hundred effective clusters after homology grouping, and a signal dominated by composition, self-attention has little long-range order-dependent structure to exploit that a composition-based representation does not already capture. The switchable components and how they compose are drawn in Supplementary Figure S4; the selected configuration retains none of them.

#### 4.5.2. Expanded search

A second optimisation round restricted to the best-performing descriptor families reached MCC 0.8345 (SD 0.054) on layer 1, against 0.8325 in the initial search, and MCC 0.8350 (SD 0.075) on layer 2, against 0.8107. The biophysical block and the SHAP-derived interaction terms were selected on both layers, and the interaction terms were the most influential search parameter on both; CatBoost was the weakest of the four classifier families (0.8053 and 0.7966). These with/without contrasts are not balanced, since the sampler concentrates on promising regions, so the direction is informative while the magnitude is not a controlled estimate. Stacking the tuned model with a length-only forest and the homology lookup, and applying isotonic or Platt calibration, both reduced out-of-fold performance on both layers. Class-imbalance handling on layer 2 under nested cross-validation produced changes within fold-to-fold variation, the best intervention yielding +0.011.

### 4.6. The mechanism recurs across peptide benchmarks

Everything above concerns one resource. Whether the mechanism is a property of this benchmark or of how peptide bioactivity benchmarks are assembled is a separate question, and it is answerable: sixteen further benchmarks are publicly available in the collection distributed with AutoPeptideML [10], spanning antibacterial, antiviral, antifungal, anticancer, ACE-[51] and DPP-IV-inhibitory, neuropeptide, toxicity and quorum-sensing activities, curated by different groups from different primary literature.

The precursor panel used in Section 4.2 does not transfer: a different bioactivity has different candidate precursors, and for most no curated panel exists. The phenomenon can nevertheless be measured without naming any precursor. If a label tracks provenance, peptides that are fragments of one another — which is what a single source protein yields — will share a label far more often than chance. We therefore group each dataset into fragment families by substring containment, exactly as in Section 4.3, and compare the proportion of label-pure multi-member families against a permutation null that preserves both the family-size distribution and the class balance. Before use, the procedure was applied to the present benchmark, where it recovers the reported 41.4% against 0.0% fragment asymmetry and returns complete family purity against a null of 0.323.

Eleven of the fourteen testable datasets in Table 5 show label purity significantly above the null after correction, nine of them at *z >* 5. Seven carry the same one-sided fragment asymmetry seen here, in which one class contains many overlapping fragments and the other almost none; DPP-IV-inhibitory peptides show the most extreme case in the collection at 63.2% against 17.8%. Three datasets carry a length confound above MCC 0.35, and one of them, T-cell antigens at 0.702, matches the 0.699 reported here.

**Table 5:**
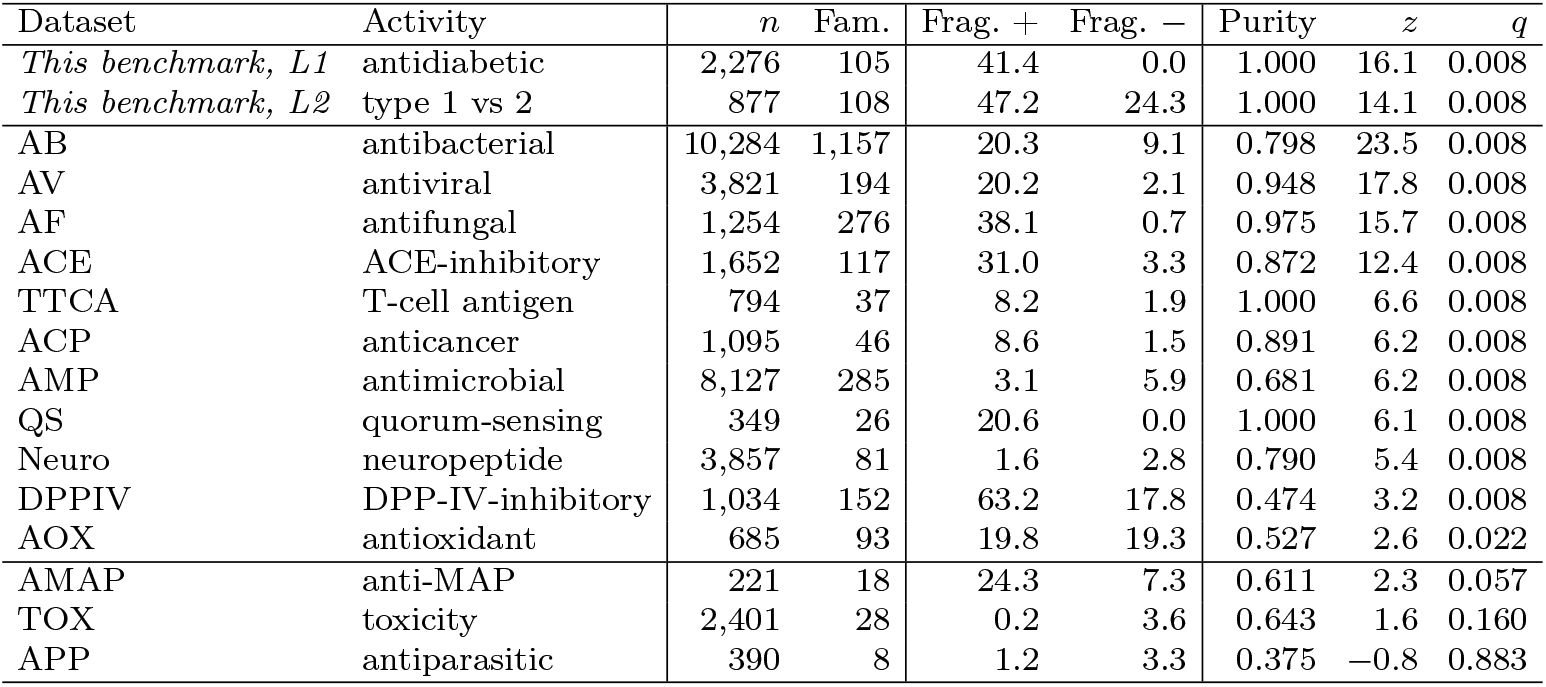
Family-label purity across sixteen further peptide benchmarks. Purity is the proportion of multimember fragment families whose members share a label; “Frag. +” and “Frag. − “are the percentages of each class that are proper fragments of another sequence. The null is a permutation preserving family sizes and class balance, and *q* is Holm-corrected across the family of tests. Two datasets, BBP (blood–brain-barrier penetrating, *n* = 191) and MRSA (*n* = 184), yielded fewer than five multi-member families and could not be tested; they are omitted rather than counted as negatives. The permutation resolution is 1*/*2000, so most *p*-values sit at the floor and *q* is correspondingly quantised; the ordering by *z* carries the information.

| Dataset | Activity | $n$ | Fam. | Frag. + | Frag. − | Purity | $z$ | $q$ |
| --- | --- | --- | --- | --- | --- | --- | --- | --- |
| *This benchmark, L1* | antidiabetic | 2,276 | 105 | 41.4 | 0.0 | 1.000 | 16.1 | 0.008 |
| *This benchmark, L2* | type 1 vs 2 | 877 | 108 | 47.2 | 24.3 | 1.000 | 14.1 | 0.008 |
| AB | antibacterial | 10,284 | 1,157 | 20.3 | 9.1 | 0.798 | 23.5 | 0.008 |
| AV | antiviral | 3,821 | 194 | 20.2 | 2.1 | 0.948 | 17.8 | 0.008 |
| AF | antifungal | 1,254 | 276 | 38.1 | 0.7 | 0.975 | 15.7 | 0.008 |
| ACE | ACE-inhibitory | 1,652 | 117 | 31.0 | 3.3 | 0.872 | 12.4 | 0.008 |
| TTCA | T-cell antigen | 794 | 37 | 8.2 | 1.9 | 1.000 | 6.6 | 0.008 |
| ACP | anticancer | 1,095 | 46 | 8.6 | 1.5 | 0.891 | 6.2 | 0.008 |
| AMP | antimicrobial | 8,127 | 285 | 3.1 | 5.9 | 0.681 | 6.2 | 0.008 |
| QS | quorum-sensing | 349 | 26 | 20.6 | 0.0 | 1.000 | 6.1 | 0.008 |
| Neuro | neuropeptide | 3,857 | 81 | 1.6 | 2.8 | 0.790 | 5.4 | 0.008 |
| DPPIV | DPP-IV-inhibitory | 1,034 | 152 | 63.2 | 17.8 | 0.474 | 3.2 | 0.008 |
| AOX | antioxidant | 685 | 93 | 19.8 | 19.3 | 0.527 | 2.6 | 0.022 |
| AMAP | anti-MAP | 221 | 18 | 24.3 | 7.3 | 0.611 | 2.3 | 0.057 |
| TOX | toxicity | 2,401 | 28 | 0.2 | 3.6 | 0.643 | 1.6 | 0.160 |
| APP | antiparasitic | 390 | 8 | 1.2 | 3.3 | 0.375 | −0.8 | 0.883 |

The three datasets below the significance threshold matter as much as the eleven above it. Antiparasitic peptides sit *below* their null, and toxicity and anti-MAP peptides do not reach significance; two further datasets had too few multi-member families to test at all. A measure that fired on every dataset would be measuring family structure rather than label–provenance coupling, and these negatives are the evidence that it does not.

What this establishes is narrower than a claim about all peptide prediction and wider than a claim about one file. Provenance-correlated labelling is a recurring property of how peptide bioactivity benchmarks are assembled — the provenance-to-function gap of the title — present in most but not all of a collection curated independently of this study, and its severity varies enough between resources that it has to be measured rather than assumed. The practical consequence is the recommendation of Section 5: report the fragment-family and precursor structure of each class when a benchmark is released, because it is cheap to compute and it determines how the reported accuracy should be read.

### 4.7. Partition sensitivity of the published design

The analysis to this point establishes that this benchmark contains structure a classifier can exploit, but it does not establish that the published figures were affected by it, because the published pipeline had not been rebuilt. That question can be asked in a weaker and answerable form: *does the published architecture lose accuracy when related peptides are prevented from straddling the split?* Answering it requires no access to the sealed independent files.

We rebuilt the multiview stacking design from its published description — 14 descriptor families, each supplying predicted probabilities and hard labels from three gradient-boosting learners to a meta-classifier — and scored it on the same cluster-disjoint folds used for our own models throughout. Meta-features for each training portion come from an inner five-fold split, so the meta-classifier never sees a base learner’s own training prediction. Table 6 reports the result.

**Table 6:** Cross-validated MCC of the rebuilt multiview design against the single tuned classifier, on identical folds and training data only. The rebuild is constructed from the published description, not from the original implementation, so it is not expected to reproduce the published values exactly; under random splitting it reaches 0.908 on layer 2 against a published 0.917, and 0.830 on layer 1 against a published 0.886, so it is a closer approximation on the second layer than on the first.

| Partition | Layer 1 |  | Layer 2 |  |
| --- | --- | --- | --- | --- |
|  | Rebuilt design | Single model | Rebuilt design | Single model |
| Random | 0.830 | 0.834 | **0.908** | 0.852 |
| Cluster-grouped $\geq 0.9$ | 0.791 | 0.826 | 0.858 | 0.837 |
| Cluster-grouped $\geq 0.7$ | 0.773 | **0.804** | 0.764 | **0.805** |
| Cluster-grouped $\geq 0.5$ | 0.778 | 0.817 | 0.812 | 0.794 |
| Loss, random $\rightarrow \geq 0.7$ | -0.057 | -0.030 | -0.144 | -0.047 |

Two things follow. First, under random splitting the rebuilt design is the stronger of the two on layer 2 (0.908 against 0.852) and not separated from it on layer 1, which is consistent with the published figures and indicates the rebuild is not a straw man. Second, that advantage does not survive homology-aware partitioning: at the primary 70% threshold the rebuilt design falls below the single model on both layers, having lost 1.9 times as much on layer 1 and 3.0 times as much on layer 2.

The interpretation must be stated carefully. This is a rebuild from a published description and not the original code, so it cannot establish what the published numbers would become under a different partition, and we do not claim that it does. What it does show is that the architecture’s advantage over a single tuned classifier is a function of how the data are split: it is present under random splitting and absent once homologues are separated. Read together with the redundancy structure of Section 4.3, this is the mechanism by which a more elaborate model can report a larger gain on this benchmark without generalising better, and it is the reason the parsimony result of Section 4.9 is not merely an argument about cost.

### 4.8. Locked independent-test evaluation

The independent test set was scored twice across this study: once following an initial lock, and once following proper tuning of the classical model family. Both are reported in Table 7. The second carries an optimistic bias of unknown magnitude relative to the first. It should be noted that no conclusion drawn in this work changes if only the first evaluation is used: layer 1 is still not separated from the published figure (0.812 against 0.841) and layer 2 remains below it (0.831 against 0.858).

**Table 7:** Locked independent-test evaluation, with both scoring events reported. ADP-Hybrid rows use the prior-corrected threshold (0.772 on layer 1, 0.262 on layer 2); all other rows use 0.5. Confidence intervals are stratified percentile bootstrap over 2,000 resamples. No confidence interval is available for the published figures, which are point estimates reproduced here exactly as reported. We note that the published first-layer sensitivity, specificity and accuracy are not mutually realisable: accuracy is a prevalence-weighted average of sensitivity and specificity and must lie between them, whereas the reported 0.940 lies below both 0.968 and 0.954 for any class balance. We compare against the published MCC because that is the metric the original study emphasises, and flag the discrepancy rather than reinterpreting their figures.

| Layer | Model | Eval. | MCC [95% CI] | ACC | Sn | Sp | AUC | PR-AUC |
| --- | --- | --- | --- | --- | --- | --- | --- | --- |
| 1 | **ADP-Hybrid** | 2 | **0.843** [0.771, 0.845] | 0.941 | 0.941 | 0.941 | 0.980 | 0.952 |
| 1 | *ADP-Fuse, as published* | – | *0.841* | *0.940* | *0.968* | *0.954* | *0.986* | – |
| 1 | Classical, all descriptors | 1 | 0.812 [0.772, 0.848] | 0.927 | 0.936 | 0.924 | 0.978 | 0.944 |
| 1 | Best neural model | 1 | 0.736 [0.689, 0.782] | 0.901 | 0.858 | 0.914 | 0.953 | 0.886 |
| 1 | Length only | 1 | 0.619 [0.570, 0.667] | 0.839 | 0.858 | 0.833 | 0.927 | 0.815 |
| 1 | Homology lookup | 1 | 0.139 [0.096, 0.178] | 0.331 | 0.963 | 0.144 | 0.809 | 0.771 |
| 2 | ADP-Hybrid | 2 | 0.801 [0.602, 0.880] | 0.955 | 0.968 | 0.864 | 0.970 | 0.995 |
| 2 | *ADP-Fuse, as published* | – | *0.858* | *0.966* | *0.968* | *0.954* | *0.993* | – |
| 2 | Locked classical | 1 | 0.831 [0.701, 0.947] | 0.960 | 0.968 | 0.909 | 0.978 | 0.997 |
| 2 | Length only | 1 | 0.633 [0.443, 0.808] | 0.927 | 0.974 | 0.591 | 0.915 | 0.983 |

### 4.9. Computational efficiency

The practical consequence of a single-model formulation is its cost. The stacked design must fit and evaluate 14 descriptor families across three gradient-boosting learners plus a meta-classifier at every inference, whereas ADP-Hybrid evaluates one classifier over one feature vector. The 14 families total 2,484 dimensions: AAC 20, DPC 400, TPC 400,

DDE 400, CTD 273, XPAAC 25, QSO 25, KSC 343, CKSAAGP 100, GXPC 30, ABC 18, BE 200, BLOS 200 and ZSC 50. Table 9 quantifies the difference on identical hardware. The stacked figures derive from a faithful rebuild of the published design rather than from the original implementation, which was unavailable during this study.

On the first layer the ADP-Hybrid point estimate is 0.843 against a published 0.841; the bootstrap interval contains the published value, and the AUC is lower (0.980 against 0.986). Because Basith et al. [5] report point estimates without confidence intervals and do not publish per-peptide predictions, no paired statistical test between the two methods is possible. We therefore state only that *these results do not distinguish the two methods*. We are not in a position to demonstrate superiority, and an unpaired interval around our own model cannot demonstrate equivalence either. On the second layer the estimate is 0.801 against 0.858 and does not exceed it. Threshold-independent behaviour for both layers is given in Supplementary Figure S8.

The second-layer independent split contains 22 negatives, and the corresponding interval spans 0.278. The consequence is quantified in Supplementary Table S33. Both methods report the same sensitivity on this layer (0.968, corresponding to 150 of 155 positives recovered), so the two reported MCC values differ only in the negative column: our model misclassifies three of the 22 negatives and the published figure two fewer. *The whole of the 0.801-versus-0.858 difference is two peptides*. Each additional false positive costs approximately 0.030 MCC, and the full range of the metric across every attainable specificity from 1.000 down to 0.727 is 0.888 to 0.709 — narrower than our own confidence interval on the same layer.

Any figure reported on this layer, whether ours or another method’s, is correspondingly uninformative. We report the shortfall because it is the result, not because the sample size excuses it; but no method comparison on this layer should be treated as evidence in either direction until the type-2 set is enlarged. We therefore regard the second-layer results throughout this paper as *hypothesis-generating rather than conclusive*, and no conclusion in Section 7 rests on them alone.

#### 4.9.1. Accuracy as a function of similarity to training data

Stratifying the locked predictions by each test peptide’s identity to the nearest training peptide (Table 8; plotted as Supplementary Figure S6) shows accuracy rising monotonically with homology. The stratum above 0.9 identity contains 129 peptides, all of which are positive, so no coefficient can be computed for it. These strata are descriptive: the lowest-identity stratum contains three positives, so no confidence intervals are attached and no inferential claim is drawn beyond the direction, which is monotone.

**Table 8:** First-layer locked-evaluation MCC stratified by each test peptide’s sequence identity to the nearest training peptide. The two arms differ only in how the model was selected before the test files were opened: the first column uses the configuration chosen on the 70%-identity cluster-grouped folds, the second the configuration chosen on the length-matched folds of Section 3.4. Both are scored on the same peptides, so the columns isolate the effect of the selection protocol rather than of the evaluation. The highest-identity stratum contains only positives and therefore cannot be scored.

| Stratum | *n* | Positives | Sel. on $\geq 0.7$ folds | Sel. on matched folds |
| --- | --- | --- | --- | --- |
| Novel, $<0.5$ | 79 | 3 | 0.428 | 0.394 |
| Remote, 0.5–0.7 | 705 | 63 | 0.719 | 0.695 |
| Similar, 0.7–0.9 | 47 | 24 | 0.918 | 0.958 |
| Near-duplicate, $\geq 0.9$ | 129 | 129 | not computable | |

**Table 9:**
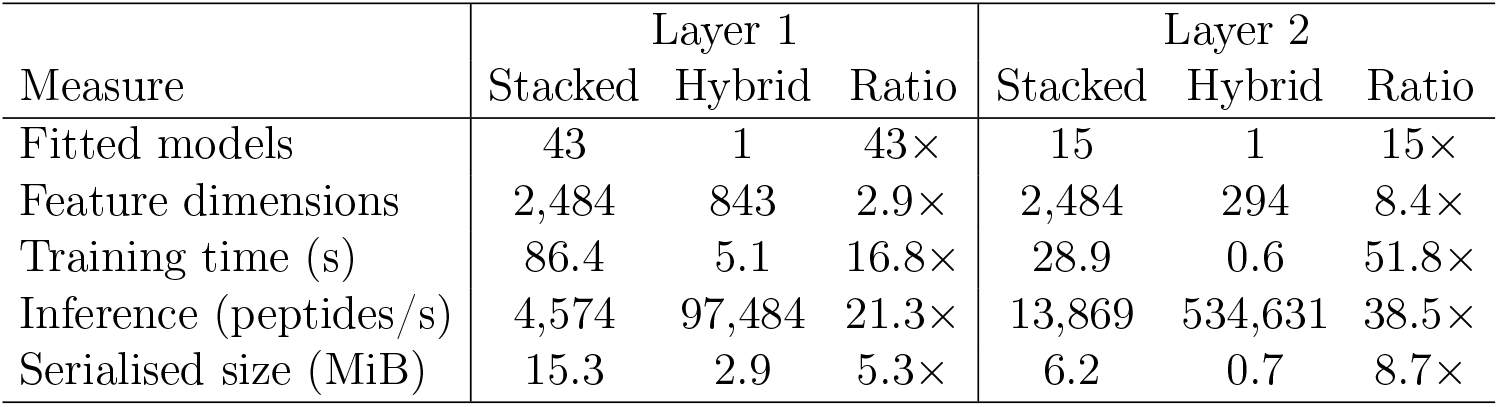
Computational cost of the stacked ensemble design against ADP-Hybrid, measured on identical hardware over the training set. **The stacked figures are obtained from a faithful rebuild of the published design — 14 descriptor families across three gradient-boosting learners plus a meta-classifier — and not from the original implementation, which was unavailable during this study. They therefore characterise the cost of that architecture as specified, not the measured cost of the released software, and an optimised implementation of the same design would fall somewhere below these figures**. Ratios are stacked relative to hybrid throughout, and are computed from unrounded measurements: a reader recomputing them from the rounded values displayed here will differ in the final digit (for example the layer-2 training ratio is 28.918/0.558 = 51.8, not 28.9/0.6 = 48.2).

### 4.10. Robustness

Seed stability was measured for the neural configuration, not for ADP-Hybrid, because it is the component with stochastic training. Five independent end-to-end retrainings under different seeds, each scored on the independent set, gave MCC 0.703 *±* 0.019 on layer 1 (*n* = 960) and 0.674 *±* 0.061 on layer 2 (*n* = 177), with AUC 0.947 *±* 0.005 and 0.958 *±* 0.017. The single best of those runs is the neural entry in Table 7 at 0.736; the spread across seeds is the reason that entry is not treated as a point estimate. ADP-Hybrid is deterministic given the frozen configuration and a fixed seed, so a seed sweep is not meaningful for it. Error analysis stratified by homology reproduces the gradient of Table 8: on layer 1 the model attains sensitivity 1.000 but specificity 0.803 in the lowest-identity stratum, against specificity 1.000 in the 0.7–0.9 stratum. Supplementary Figure S2 shows the same breakdown across homology, length and threshold axes together.

Stratifying instead by peptide length (Supplementary Table S25) exposes a second single-class structure. On the second layer the two longest quartiles, 90 of the 177 independent-test peptides, contain no type-2 peptide at all: every test peptide longer than 15 residues is type-1. No coefficient exists for those strata, and they are reported as such rather than dropped. The first layer shows the same effect in weaker form, its longest quartile holding one positive among 248 peptides. Length and class are thus confounded in both layers of this benchmark, which is the partition-level counterpart of the length result in Section 4.4.

## 5. Discussion

The claims this study makes, and the claims it deliberately does not make, are set out point by point in Supplementary Section S14; the limitations of Section 6 state the same boundaries in prose.

### What this benchmark measures

The dominant exploitable signal in this dataset is closely related to provenance. The first layer separates a collection of peptide-hormone fragments from a collection of antimicrobial and cell-penetrating peptides that differ from them by roughly twelve residues in mean length and substantially in composition. The second layer separates human-insulin fragments from bovine milk-protein fragments, at least within the subset that could be attributed. A model may therefore score highly on both layers without having learned a determinant of antidiabetic activity as such.

### Generality

The mechanism is not specific to this resource. Curated activity databases accrete entries study by study, a study characterising a single precursor contributes many overlapping fragments at once, and a label defined by clinical indication will track provenance whenever different indications were historically investigated in different source proteins. Section 4.6 shows that this is not merely plausible: label–provenance coupling is significant in eleven of fourteen independently curated peptide benchmarks. What remains bounded is the precursor-level demonstration, which required a curated panel and was therefore possible only here; elsewhere the coupling is established through family structure rather than named source proteins.

### Practical contribution

A single classifier over 843 features reaches first-layer accuracy the evidence does not separate from the ensemble, at 21–38*×* the inference throughput and a fifth to a ninth of the footprint, and Section 4.7 shows it also degrades less once homologues are separated. The saving is therefore not only in cost: where a 43-model ensemble must be maintained in its entirety, a single model can be inspected feature by feature.

### Why additional capacity did not help

Across two optimisation rounds, six architectures, stacking, calibration and class-imbalance handling, nothing exceeded a tuned single classifier by a margin exceeding fold-to-fold variation. Read together with the length and homology results, the natural interpretation is that the accessible signal is largely low-order and composition-like, and had already been saturated.

### Recommendations for future peptide-function benchmarks

Five measures follow. Apply redundancy reduction symmetrically to both classes, since applying it to one creates a class-dependent difficulty gradient. Report each class’s fragment-family and precursor structure, which Section 4.6 shows is cheap to compute and informative in most benchmarks. Report a length-only baseline, which costs one line of code and bounds how much of a result a trivial covariate explains. Report accuracy stratified by identity to the training set, flagging single-class strata. Partition multi-stage predictors once across all stages, so the deployed cascade has an honest end-to-end estimate.

## 6. Limitations

1. The independent test set was scored twice. Both evaluations are reported; the second carries an optimistic bias of unknown magnitude. No conclusion changes under the first evaluation alone.
2. No paired significance test against ADP-Fuse was possible, as point estimates are published without confidence intervals, per-peptide predictions are not available, and the associated web server did not respond during this study. The intervals reported here are unpaired estimates of our own model’s performance. Consequently neither superiority nor equivalence is claimed on layer 1.
3. The published design was rebuilt from its description and shown to be more partition-sensitive than a single tuned model (Section 4.7), but the rebuild is not the original implementation. What the published *figures* would become under a homology-aware partition therefore remains untested, and nothing here establishes that they are inflated.
4. Precursor attribution covers 24.9% of the set, is a lower bound, and the attributed subset is not a random sample of the whole.
5. The second layer cannot separate methods at 22 independent-test negatives.
6. Architecture conclusions are bounded to matched conditions on this dataset, with frozen ESM-2 representations and a 60-trial neural search budget.
7. The precursor-level attribution — naming the source protein for each peptide — was possible only for this benchmark, because it required a curated UniProt panel. Across the sixteen further benchmarks the coupling is established through fragment-family structure alone, which shows that a label tracks provenance without identifying what that provenance is.
8. Three of the fourteen testable benchmarks show no significant coupling and two more could not be tested for want of multi-member families, so the phenomenon is common rather than universal and has to be measured per resource.

## 7. Conclusion

Within the subset of an antidiabetic peptide benchmark whose peptides can be attributed to a precursor protein, the second-layer class label is fully determined by that precursor. Predictive accuracy on the benchmark tracks sequence identity to the training set, and peptide length alone recovers a substantial share of the reported performance. Against this background, a single tree-ensemble classifier reaches first-layer accuracy that the available evidence does not separate from a published 43-model ensemble, at a fraction of the computational cost, and did not exceed it on the second layer. These are offered as constructive observations concerning a resource that has been valuable to the field, together with an evaluation protocol and a set of controls intended to assist in constructing the next generation of peptide-function benchmarks.

## Data and code availability

All analysis code, the corrected evaluation protocol, the frozen model configuration and every results file listed in Supplementary Table S32 are available from the corresponding author on reasonable request.

## Supplementary Information

One file, containing 34 tables and 7 figures across fourteen sections, each with explanatory text; its contents are listed on the first page of that file. Table S32 maps every claim in this manuscript to the script that produces it and the results file that records it.

Precursor sequences were retrieved from UniProt [18]. The benchmark data are those distributed by Basith et al. [5] and are not redistributed here. Supplementary Table S32 lists what is released against each claim in this paper.

## Acknowledgements

We thank the authors of ADP-Fuse [5] for curating and publicly distributing the two-layer benchmark and web server that made this study possible, and the authors of BioDADPep [3] for the underlying resource.

